# Population-level temporal decoding and dynamical structure in frontostriatal circuits during decision making

**DOI:** 10.1101/2025.11.07.687310

**Authors:** Adam B. Goldring, Anandita De, Tanner Stevenson, Kendall Stewart, Aya Akhmetzhanova, Rishidev Chaudhuri, Timothy D. Hanks

## Abstract

Perceptual decisions unfold over time, requiring neural circuits to evaluate sensory evidence, track elapsed time, and commit to an action. To investigate how these computations are distributed across corticostriatal circuits, we recorded large-scale neural population activity using Neuropixels probes in the rat frontal orienting field (FOF) and anterior dorsal striatum (ADS) during a free-response auditory change detection task. Both regions exhibited structured, highly dynamic population activity during task performance. Using single-trial population decoding, we found that both FOF and ADS robustly encoded retrospective time from stimulus onset and prospective time preceding the decision report. Using a new approach to analyze temporal encoding to identify the dynamical structure supporting decoding, we found that time encoding could be decomposed into two primary dynamical motifs: monotonic ramp-like trajectories and transient bump-like trajectories. While these modes were similarly expressed during early evidence evaluation, FOF exhibited more pronounced decision-aligned changes near the time of the decision report, possibly reflecting a state transition at commitment. Population geometry analyses further revealed stable low-dimensional subspaces during evidence evaluation that transitioned at decision commitment, with significantly larger subspace changes in FOF than ADS. Together, these results reveal a common dynamical framework during evidence evaluation across corticostriatal circuits, while identifying a selective reorganization of FOF dynamics associated with decision commitment.

## Introduction

Adaptive behavior requires organisms to make decisions based on sensory evidence. In many situations, relevant evidence must be evaluated sequentially, so the decision process unfolds over time. A long literature has identified brain regions that are involved in evidence evaluation and decision commitment, particularly across frontal cortex and striatal circuits (Brody and Hanks, 2016; Ding and Gold, 2013; Gold and Shadlen, 2007; Hanks et al., 2015; Hanks and Summerfield, 2017; Khilkevich et al., 2024; Yartsev et al., 2018). It is therefore important to understand the population-level temporal dynamics of these distributed circuits during decision making.

The frontal cortex and striatum have long been implicated as critical brain regions for both temporal processing and decision making (Affan et al., 2025; Emmons et al., 2017; Gold and Shadlen, 2007; Jin et al., 2009; Kim et al., 2013; Matell and Meck, 2004; Mauk and Buonomano, 2004; Merchant et al., 2013; Murakami et al., 2017, 2014; Sosa et al., 2021; Zhou et al., 2020). In rodents, the highly interconnected frontal orienting field (FOF) and anterior dorsal striatum (ADS) have been identified as key nodes in a decision making network involved in evidence evaluation and decision commitment (Brody and Hanks, 2016; Ding and Gold, 2013; Erlich et al., 2011; Hanks et al., 2015; Yartsev et al., 2018). Past work suggests important overlap and some possible distinctions in contributions of these regions (DePasquale et al., 2024; Hanks et al., 2015; Yartsev et al., 2018; Zhou et al., 2020). However, large-scale simultaneous recordings of neurons from both areas are needed to better understand whether their contributions primarily reflect shared computations, or whether they diverge in function at specific phases of the decision process.

One dimension along which these circuits may differ or interact is in their encoding of time. Knowledge of elapsed or remaining time benefits many components of decision making, such as monitoring how long evidence has been presented, recognizing temporal patterns of evidence, or anticipating possible changes in evidence. Time representations have been observed in both the frontal cortex and striatum (Gouvêa et al., 2015; Jin et al., 2009; Kim et al., 2013; Mello et al., 2015; Wang et al., 2018; Xu et al., 2014; Zhou et al., 2020). However, whether temporal representations during decision making in frontal and striatal circuits share common dynamical structure, and how they reorganize during the transition from evidence evaluation to decision commitment, remains largely unknown.

Population-level analyses of large-scale neural recordings offer a way to address these questions. Neural circuits supporting decision making may contain diverse and heterogeneous single-neuron response patterns, yet exhibit low-dimensional trajectories at the population level that reveal underlying dynamical structure. Characterizing these population trajectories and how they reorganize across task epochs can identify the extent to which circuits share computational roles or differ.

Here, we leverage approaches that combine single-trial decoding from population responses and analysis of population geometry to investigate temporal encoding and population dynamics in rat FOF and ADS during a free-response auditory change detection task. This task requires rats to monitor a stochastic auditory stream, detect a change in the stimulus statistics, and initiate a behavioral response at the time of their choice to report changes. Using Neuropixels recordings, we quantified how neural populations in FOF and ADS encoded retrospective elapsed time and prospective time to the choice on individual trials, and we developed an analytical framework to identify dynamical motifs that support temporal decoding. We find that both regions exhibit two matched temporal modes: a monotonic ramp and a transient bump. Differences between the regions emerge near the time of decision commitment. By tracking the geometry of population activity across task epochs, we show that while both regions maintain stable subspaces during evidence evaluation, the FOF undergoes a significantly larger change in subspace at the time of decision commitment. Together, these results reveal both shared and divergent temporal dynamics across frontal and striatal circuits, suggesting that while FOF and ADS jointly support temporal processing during decision making, FOF exhibits a distinct shift in population state at commitment.

## Results

### Neural recordings and behavior

We used Neuropixels probes to record neural spiking activity from FOF and ADS while rats performed a free-response auditory change detection task (**Figure 1a-c**). Our analyses focus on population-level activity within single trials and single sessions, so we included both single units and multi-unit clusters. In total, we analyzed 1523 units from FOF (814 single and 709 multi) and 7235 units from ADS (4662 single and 2573 multi).

**Figure 1.**
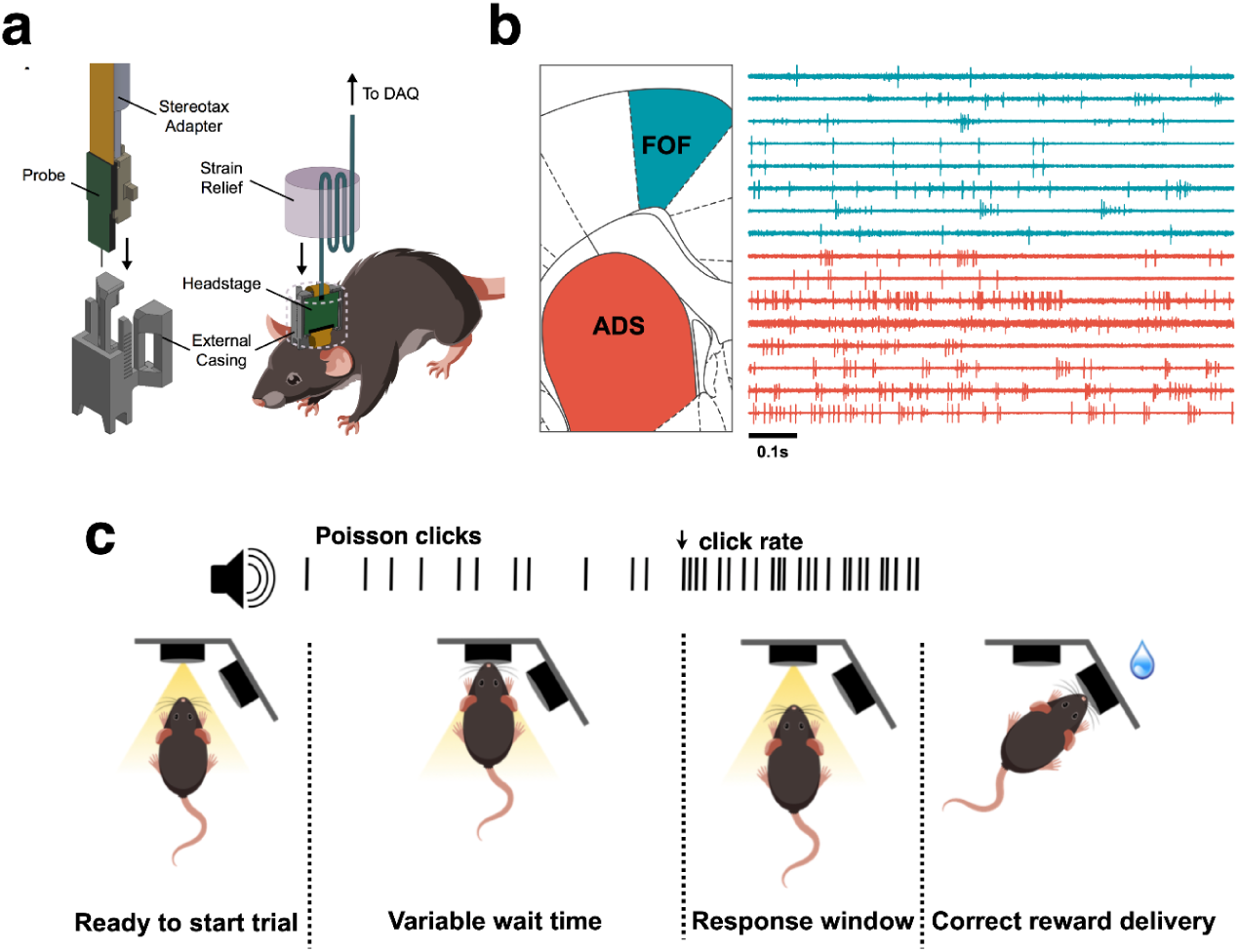
Experimental approach. **a**, For each rat, a Neuropixels recording probe was implanted using a custom designed assembly that allowed use of a strain-relieving attachment for the tether. **b**, The probe was positioned to record simultaneously from both FOF and ADS, with the schematic showing the coronal plane. The right side shows traces from example sites recorded across both regions simultaneously. **c**, Task schematic.

In the task, rats are cued with a light to insert their noses into a nose port of an operant apparatus, which initiates a stream of broadband auditory pulses (“clicks”) generated via a Poisson process. Starting at a baseline rate that is fixed for each session, the generative click rate may increase by a variable magnitude at a random time at least 500 ms after stimulus onset, to which subjects must respond within a fixed window by withdrawing their nose from the port. A timely withdrawal in response to a change is a *hit*, which leads to a water reward delivered from a separate port on the side of the apparatus. Failure to withdraw in time results in a *miss*, with no reward. Withdrawal in the absence of a change is a *false alarm*, also denying a reward. Finally, on a subset of trials, no change in stimulus rate occurs; during these “catch” trials, the rat must maintain port fixation until the stimulus ends, resulting in a *correct rejection* accompanied by a water reward. The task has been used previously in rats (Ganupuru et al., 2025) and was adapted from previous studies involving human subjects (Johnson et al., 2017).

Similar to previous studies, rats consistently detected changes, achieving higher hit rates with higher change magnitudes (Regression coefficient: 0.086 +/- 0.008, p < 0.001; **Figure 2a**), although they were prone to false alarms (average false alarm rate was 0.68 +/- 0.01). Higher change magnitudes also yielded faster reaction times (Regression coefficient: -2.73 +/- 0.30, p < 0.001; **Figure 2b**). These metrics suggest rats perform the task by evaluating the presented sensory evidence, and greater quantities of evidence lead to faster and more accurate choices. To better gauge the strategy rats employ on this task, we conducted psychophysical reverse correlation (PRC) analyses using false alarm trials to assess how rats evaluated evidence over time prior to their decision to withdraw their nose from the port. PRC traces were constructed by convolving click times with causal half-Gaussian filters (sigma = 0.05 s) and aligning the result to the time of the false alarm response. Because the Poisson stimulus varies around a generative mean from moment to moment, this analysis reveals how false alarms are related to stochastic fluctuations in click rate as a function of time. Therefore, the PRC method provides insight into the timescale and strength of evidence contributing to decision formation (Ganupuru et al., 2019; Okazawa et al., 2018).

**Figure 2.**
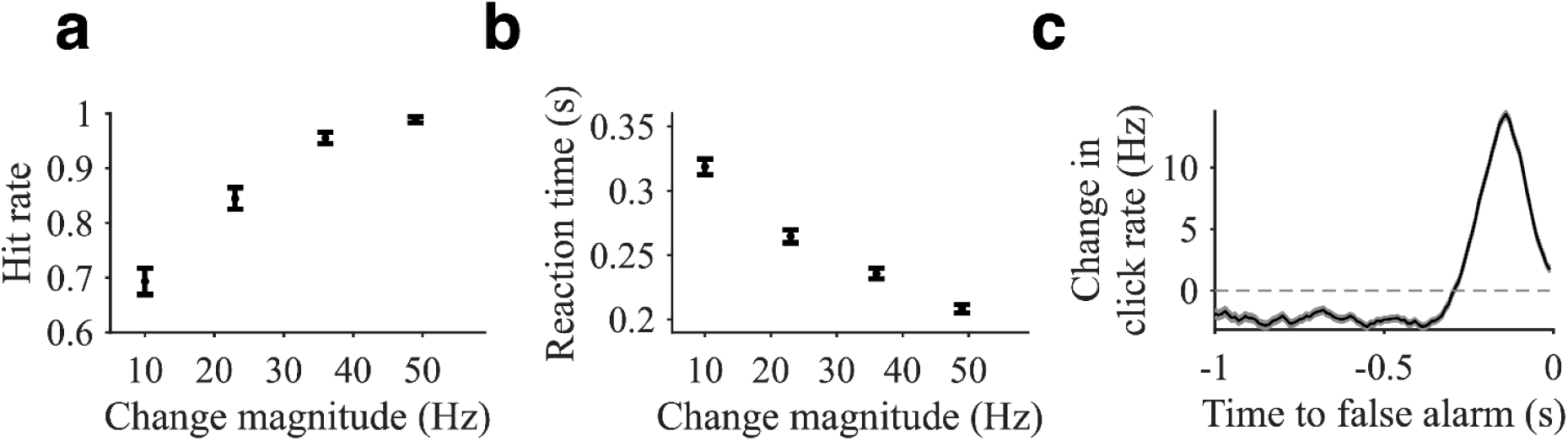
Behavioral performance. **a**, Combined hit rates as a function of change magnitude. Hit rate was calculated excluding false alarm trials (i.e. only including trials in which the rat was presented with a change). **b**, Reaction times for hits measured as the time of center port withdrawal following a change. **c**, Psychophysical reverse correlation (PRC). Calculated as the average local click rate preceding false alarms. For all plots, error bars and error shading show SE.

As expected, the false alarm PRC exhibited an upward fluctuation in click rate prior to false alarms, indicating false alarms were generally associated with brief increases in click rate (**Figure 2c**). The mean increase of the click rate from 50 to 350 ms before the false alarm was 7.2 +/- 0.1 Hz (p < 0.001). The PRC thus suggests a behavioral strategy on the part of the rats similar to previous studies (Ganupuru et al., 2025).

### Temporal Decoding

The auditory change detection task described above requires the rats to perform several related but distinct operations in order to successfully complete a trial. Many of these operations may involve and benefit from the tracking of elapsed time. This includes determining when the 500 ms minimum pre-change period has ended, learning the duration of the response window, and estimating click rates over differing temporal intervals. Therefore, the first-order question we sought to answer is the degree to which neuronal responses in these regions can predict temporal interval relative to specific behaviorally relevant events (e.g., start of the stimulus, movement initiation, etc.).

We addressed this question by performing temporal decoding at a single-trial level based on spiking responses from populations of neurons recorded simultaneously in FOF and ADS. These analyses were performed separately for individual behavior sessions (N = 61 sessions in total). We first sought to decode time elapsed from stimulus start. To ensure that no extrinsic factors beyond time influenced this decoding, we focused on trials that had an uninterrupted period of at least 1 full second from stimulus onset until change in the generative click rate or movement response initiation. We used a “leave one trial out” decoding procedure where performance of the decoder in predicting time within that 1 second period for each trial was based on training the decoder on all other trials except the one being decoded, and this was done iteratively for every trial. **Figure 3a** and **3b** show decoding performance for an example session for neurons recorded in ADS and FOF, respectively. Predicted times were generally in close alignment with the true time from stimulus onset, as illustrated by the heavy weight clustered around the diagonal.

**Figure 3.**
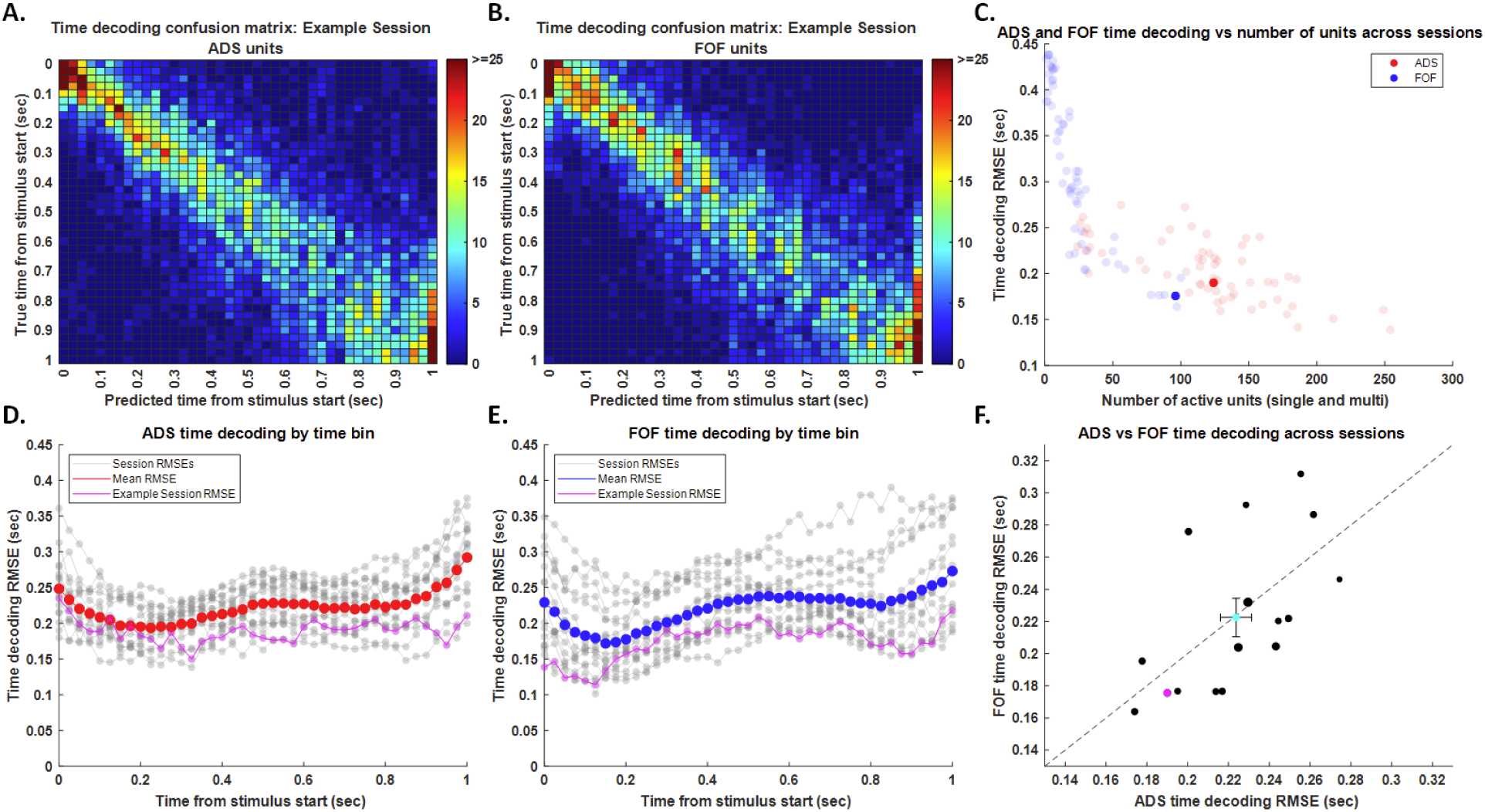
Retrospective temporal decoding from stimulus onset. **a**, Example decoding “confusion matrix” derived from ADS units. Color bar indicates number of trials (out of 169) classified into each bin comparing predicted to true time from stimulus onset. **b**, Decoding confusion matrix from FOF units in the same session. **c**, Average root-mean squared error (RMSE) of temporal decoding for each session as a function of the total number of units in that session, separated by region. Darker points correspond to the example session shown in earlier panels. N = 61 sessions. **d**, ADS temporal decoding RMSE as a function of time for each analyzed session (gray traces). Magenta trace shows example session from earlier panels. Red trace shows average RMSE across sessions. **e**, FOF temporal decoding RMSE as a function of time for each analyzed session (gray traces). Magenta trace shows example session from earlier panels. Blue trace shows average RMSE across sessions. **f**, Session by session comparison of RMSE for FOF versus ADS. Magenta point shows example session from earlier panels. Cyan point shows mean and SEM across sessions. Only sessions that meet criteria for balance between ADS and FOF units are included in panels **d-f**. N = 16 sessions.

In comparing decoding accuracy across brain regions, it is important to consider the impact of the total number of units used for the decoding. Decoding accuracy was quantified as the root mean squared error (RMSE) in seconds between the predicted and true time. For both regions, there was a clear and significant trend for improved decoding accuracy with higher numbers of units (ADS slope: -4.3e-04 +/- 6e-05, p < 0.001; FOF slope -2.8e-03 +/- 3e-04, p < 0.001; **Figure 3c**).

Because recordings were performed simultaneously in ADS and FOF, we can perform direct session-by-session comparisons of decoding accuracy. For these comparisons, we excluded sessions with highly unbalanced unit counts (fewer than 40% in one area compared to the other) and instead focused on the remaining 16 sessions. We first examined whether either region’s decoding accuracy depended on time (**Figure 3d,e**). For both regions, we found fluctuations in decoding accuracy as a function of time from stimulus start with a significant trend for worsening as time elapsed (ADS slope: 0.045 +/- 0.009, p < 0.001; FOF slope 0.066 +/- 0.008, p < 0.001). Collapsing across time, we found that the RMSE for these sessions was comparable for ADS and FOF (224 +/- 8 ms for ADS; 223 +/- 12 ms for FOF; p = 0.90 for difference; **Figure 3f**).

The previous analyses used firing rates to decode retrospective time elapsed from stimulus onset in the past. A similar approach can be used to decode prospective time until movement initiation in the future. **Figure 4a** and **4b** show prospective time decoding performance for the same example session as illustrated above for retrospective decoding performance. Less accurate decoding performance for prospective time leads to more scattered predictions and less weight clustered around the diagonal than in the same session for retrospective time decoding. Similar to retrospective time decoding, there was a significant trend for improved prospective time decoding accuracy with higher numbers of units in both areas (ADS slope: -2.0e-04 +/-6e-05, p = 0.002; FOF slope -2.3e-03 +/- 2e-04, p < 0.001; **Figure 4c**). Also similar to retrospective time decoding, we found a significant change in prospective decoding accuracy as a function of time but with improved accuracy as time elapsed (ADS slope: -0.055 +/- 0.017, p = 0.002; FOF slope -0.12 +/- 0.01, p < 0.001; **Figure 4d,e**). This makes intuitive sense because in both cases, accuracy was better closer to “anchor times” (stimulus start for retrospective time decoding and movement initiation for prospective time decoding). Interestingly, collapsing across time, we found that the average RMSE was lower for prospective time decoding from FOF compared to ADS (340 +/- 9 ms for ADS; 321 +/- 11 ms for FOF; p = 0.008 for difference; **Figure 4f**), even though there were generally fewer units recorded per session in FOF. This effect was significant in the interval from 525 to 0 ms before the movement, suggesting an emerging difference in FOF versus ADS nearing the time of the decision report.

**Figure 4.**
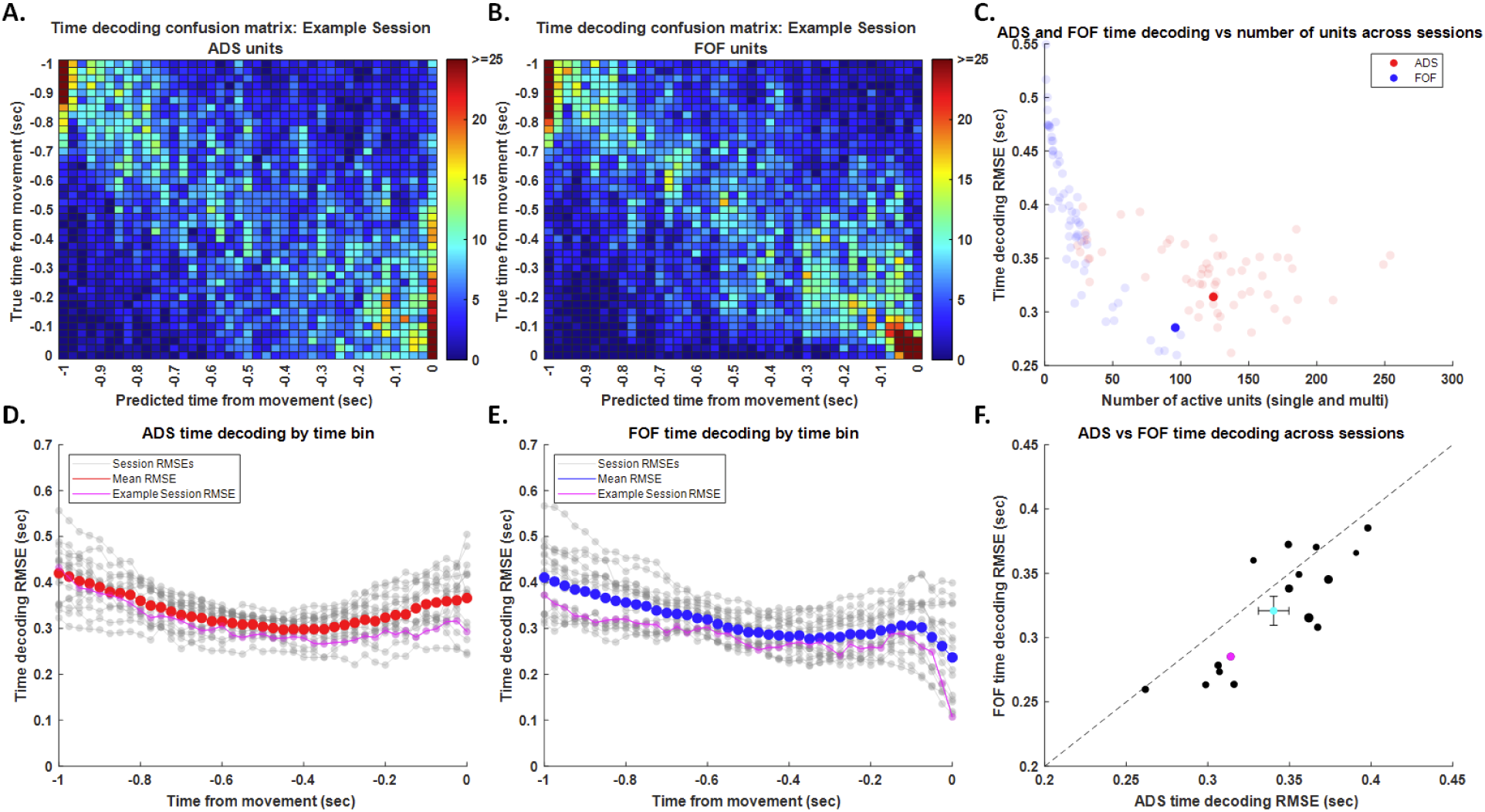
Prospective temporal decoding to movement initiation. **a**, Example decoding “confusion matrix” derived from ADS units. Color bar indicates number of trials (out of 214) classified into each bin comparing predicted to true time from stimulus onset. **b**, Decoding confusion matrix from FOF units in the same session. **c**, Average root-mean squared error (RMSE) of temporal decoding for each session as a function of the total number of units in that session separated by region. Darker points correspond to the example session shown in earlier panels. N = 61 sessions. **d**, ADS temporal decoding RMSE as a function of time for each analyzed session (gray traces). Magenta trace shows example session from earlier panels. Red trace shows average RMSE across sessions. **e**, FOF temporal decoding RMSE as a function of time for each analyzed session (gray traces). Magenta trace shows example session from earlier panels. Blue trace shows average RMSE across sessions. **f**, Session by session comparison of RMSE for FOF versus ADS. Magenta point shows example session from earlier panels. Cyan point shows mean and SEM across sessions. Only sessions that meet criteria for balance between ADS and FOF units are included in panels **d-f**. N = 16 sessions.

### Neural dynamics

Given the capacity to decode both retrospective and prospective time within individual trials from firing rates in each region, we next sought to understand the temporal dynamics of neural activity in each region that underpin time decoding, and more fundamentally, how the manner in which neurons contribute to time decoding relates to distinctive firing rate dynamics. To examine this question, we developed an analytical framework to sort units based on their contributions to temporal decoding. We started with retrospective temporal decoding from stimulus onset and applied this analysis separately to units from ADS and FOF. The temporal decoder results in a set of weights reflecting the contribution of each unit to classification as a function of time, with individual units potentially having distinct patterns of contributions. Principal components analysis (PCA) of the unit-by-time weight matrices was used to capture these patterns in a low-dimensional space (**Figure 5**,**6a**). Sorting units based on their positions within this space provides a population-level view of how diverse neuronal dynamics can support temporal decoding. In the trivial case of no sorting (**Figure 5**,**6a**), the overall population neural dynamics were flat for both ADS and FOF(**Figure 5**,**6b-c**). This suggests that if there are diverse temporal dynamics between neurons, they largely balance each other out when averaged together. More interesting dynamics were revealed when units were sorted based on the first principal component (**Figure 5**,**6d**). For this case, we found that variation in that dimension revealed differences in steepness of ramping dynamics for both ADS and FOF (**Figure 5**,**6e-f**). When units were sorted based on the second principal component (**Figure 5**,**6g**), rather than ramping dynamics, it revealed more transient differences that did not persist and tended to collapse at the end of the temporal interval (**Figure 5**,**6h-i**). This suggests a common feature between ADS and FOF of two distinct dynamical modes, one more persistent and one more transient, that underlie temporal decoding. Sorting units based on both principal components (**Figure 5**,**6j**) allows for composition of these two modes that can result in more complex dynamics including delayed ramping and early ramping followed by a plateau (**Figure 5**,**6k-l**).

**Figure 5.**
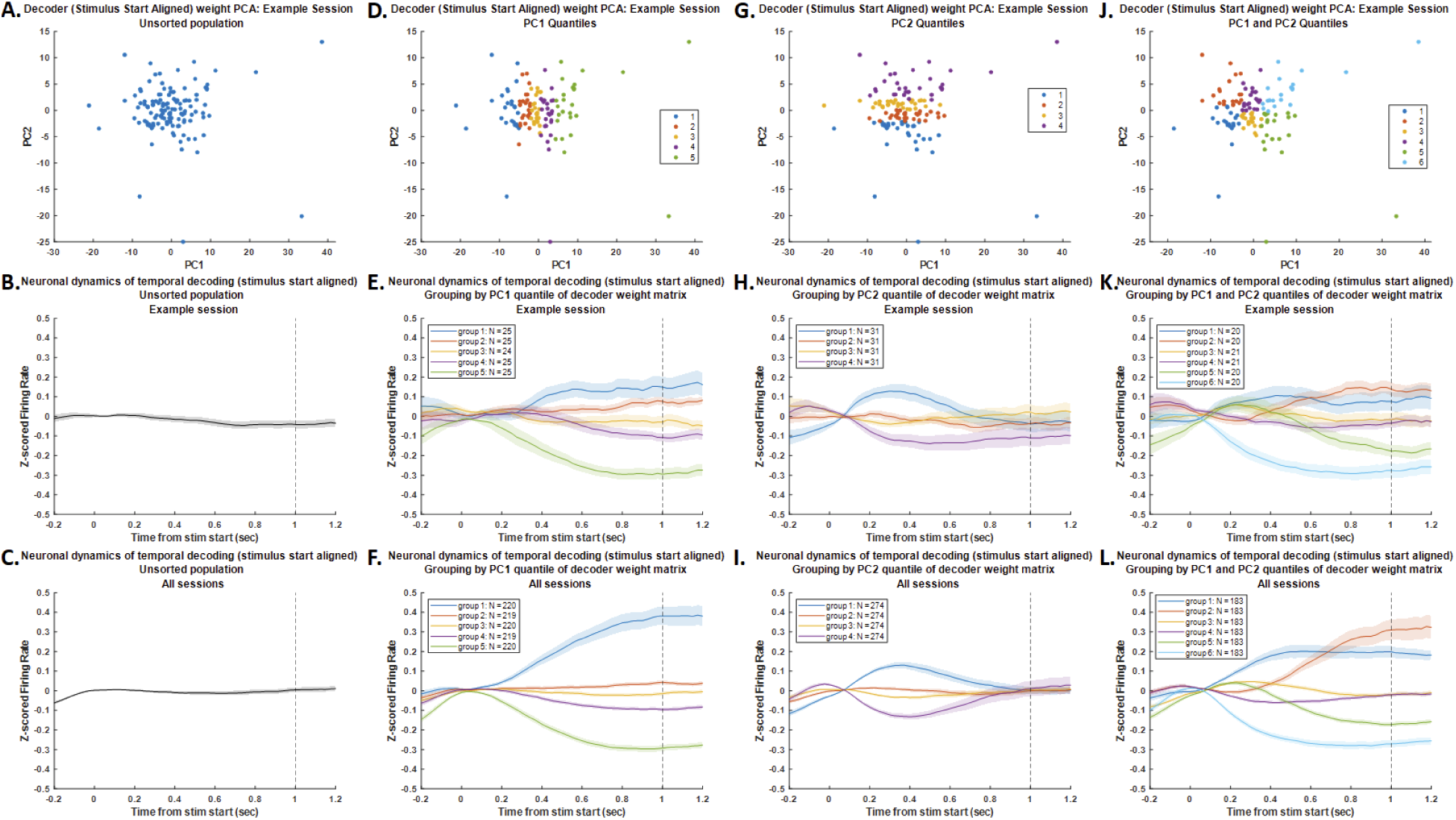
Retrospective temporal encoding from stimulus onset in ADS. **a**, Example session showing each unit plotted in the space of the first two principal components of the decoder weights. Each point corresponds to one unit. **b**, z-scored firing rate of all units from this session aligned to stimulus start. The vertical line corresponds to the end of the decoding epoch for all panels showing neural dynamics. **c**, z-scored firing rate of all units from all sessions aligned to stimulus start. **d**, Same as panel **a**, but with units sorted into quintiles along the first principal component. **e**, z-scored firing rate of all units from this session sorted into quintiles along the first principal component and aligned to stimulus start. **f**, z-scored firing rate of all units from all sessions sorted into quintiles along the first principal component and aligned to stimulus start. **g**, Same as panel **a**, but with units sorted into quartiles along the second principal component. **h**, z-scored firing rate of all units from this session sorted into quartiles along the second principal component and aligned to stimulus start. **i**, z-scored firing rate of all units from all sessions sorted into quartiles along the second principal component and aligned to stimulus start. **j**, Same as panel **a**, but with units sorted into six groups along both of the first two principal components. **k**, z-scored firing rate of all units from this session sorted into six groups along both of the first two principal components and aligned to stimulus start. **l**, z-scored firing rate of all units from all sessions sorted into six groups along both of the first two principal components and aligned to stimulus start. N = 16 sessions.

**Figure 6.**
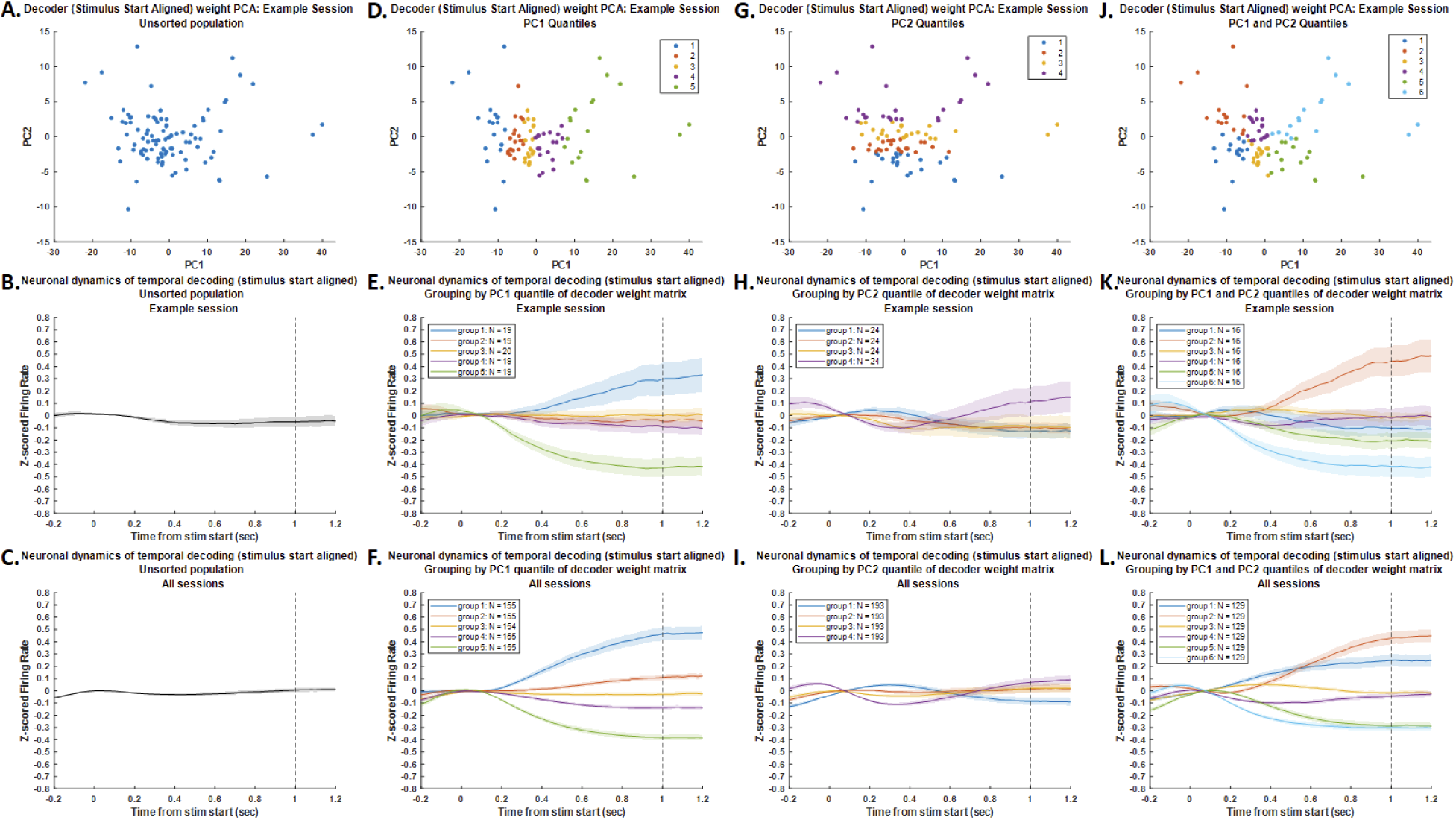
Retrospective temporal encoding from stimulus onset in FOF. **a**, Example session showing each unit plotted in the space of the first two principal components of the decoder weights. Each point corresponds to one unit. **b**, z-scored firing rate of all units from this session aligned to stimulus start. The vertical line corresponds to the end of the decoding epoch for all panels showing neural dynamics. **c**, z-scored firing rate of all units from all sessions aligned to stimulus start. **d**, Same as panel **a**, but with units sorted into quintiles along the first principal component. **e**, z-scored firing rate of all units from this session sorted into quintiles along the first principal component and aligned to stimulus start. **f**, z-scored firing rate of all units from all sessions sorted into quintiles along the first principal component and aligned to stimulus start. **g**, Same as panel **a**, but with units sorted into quartiles along the second principal component. **h**, z-scored firing rate of all units from this session sorted into quartiles along the second principal component and aligned to stimulus start. **i**, z-scored firing rate of all units from all sessions sorted into quartiles along the second principal component and aligned to stimulus start. **j**, Same as panel **a**, but with units sorted into six groups along both of the first two principal components. **k**, z-scored firing rate of all units from this session sorted into six groups along both of the first two principal components and aligned to stimulus start. **l**, z-scored firing rate of all units from all sessions sorted into six groups along both of the first two principal components and aligned to stimulus start. N = 16 sessions.

While differences in dynamics aligned to stimulus onset revealed by this analysis between ADS and FOF were subtle at most, clearer distinctions did emerge when applying the same framework to prospective decoding of time before movement initiation. In the trivial case of no sorting (**Figure 7**,**8a**), the overall population neural dynamics aligned to movement initiation were flat for both ADS and FOF (**Figure 7**,**8b-c**). When units were sorted based on the first principal component (**Figure 7**,**8d**), there were also overall ramping dynamics (**Figure 7**,**8e-f**). However, there appeared to be a large movement-aligned increase in the most extreme subset of FOF units that is not present in ADS (compare group 5 in **Figure 7**,**8e-f**). When units were sorted based on the second principal component (**Figure 7**,**8g**), ADS appeared to have more subtle differences than FOF (**Figure 7**,**8h-i**). This was particularly apparent when sorting units based on both principal components (**Figure 7**,**8j**), where ADS dynamics followed relatively steady trajectories while FOF dynamics appeared to change more dramatically near the time of movement initiation (**Figure 7**,**8k-l**).

**Figure 7.**
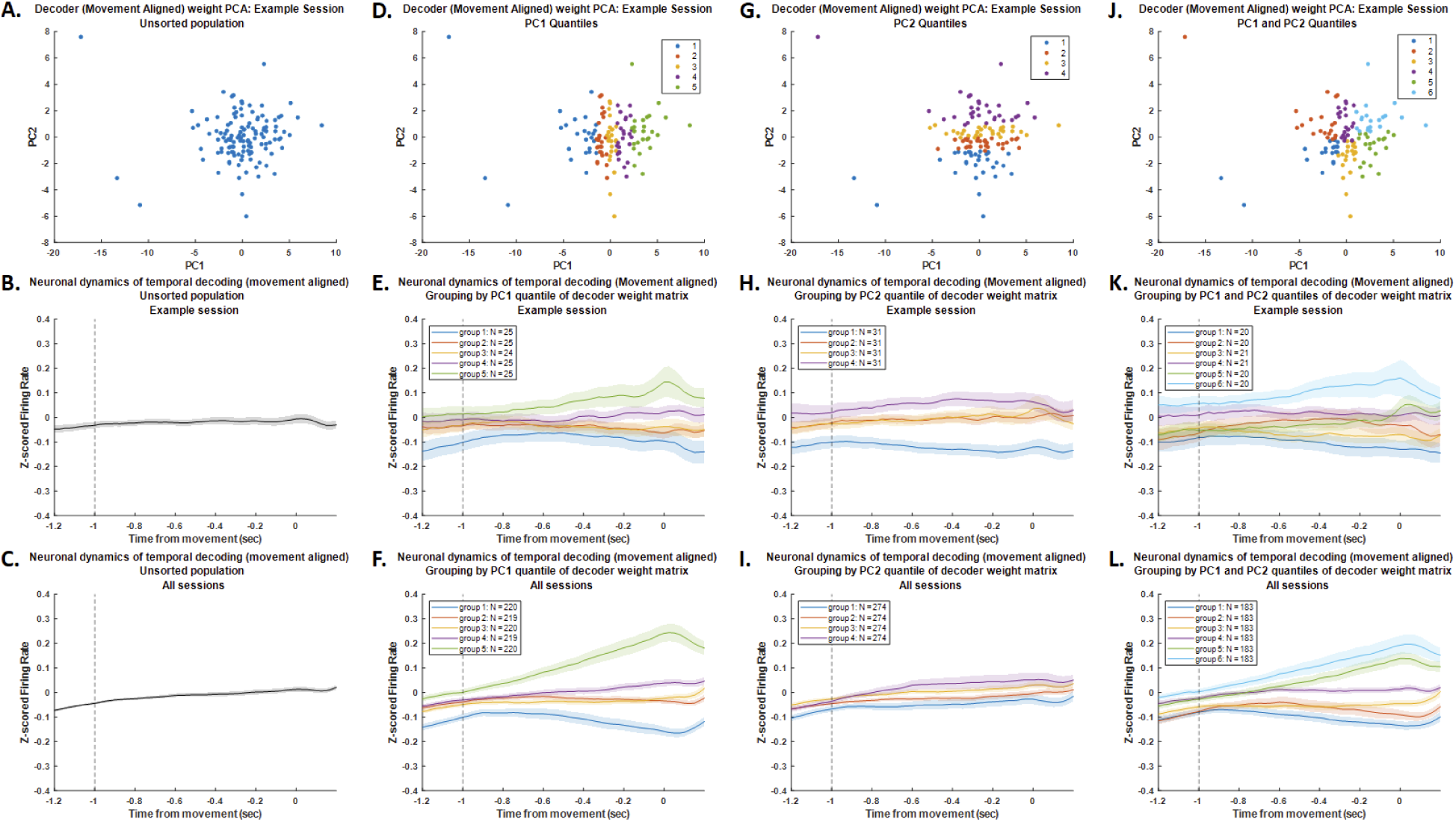
Prospective temporal encoding to movement initiation in ADS. **a**, Example session showing each unit plotted in the space of the first two principal components of the decoder weights. Each point corresponds to one unit. **b**, z-scored firing rate of all units from this session aligned to movement onset. The vertical line corresponds to the end of the decoding epoch for all panels showing neural dynamics. **c**, z-scored firing rate of all units from all sessions aligned to movement onset. **d**, Same as panel **a**, but with units sorted into quintiles along the first principal component. **e**, z-scored firing rate of all units from this session sorted into quintiles along the first principal component and aligned to movement onset. **f**, z-scored firing rate of all units from all sessions sorted into quintiles along the first principal component and aligned to movement onset. **g**, Same as panel **a**, but with units sorted into quartiles along the second principal component. **h**, z-scored firing rate of all units from this session sorted into quartiles along the second principal component and aligned to movement onset. **i**, z-scored firing rate of all units from all sessions sorted into quartiles along the second principal component and aligned to movement onset. **j**, Same as panel **a**, but with units sorted into six groups along both of the first two principal components. **k**, z-scored firing rate of all units from this session sorted into six groups along both of the first two principal components and aligned to movement onset. **l**, z-scored firing rate of all units from all sessions sorted into six groups along both of the first two principal components and aligned to movement onset. N = 16 sessions.

**Figure 8.**
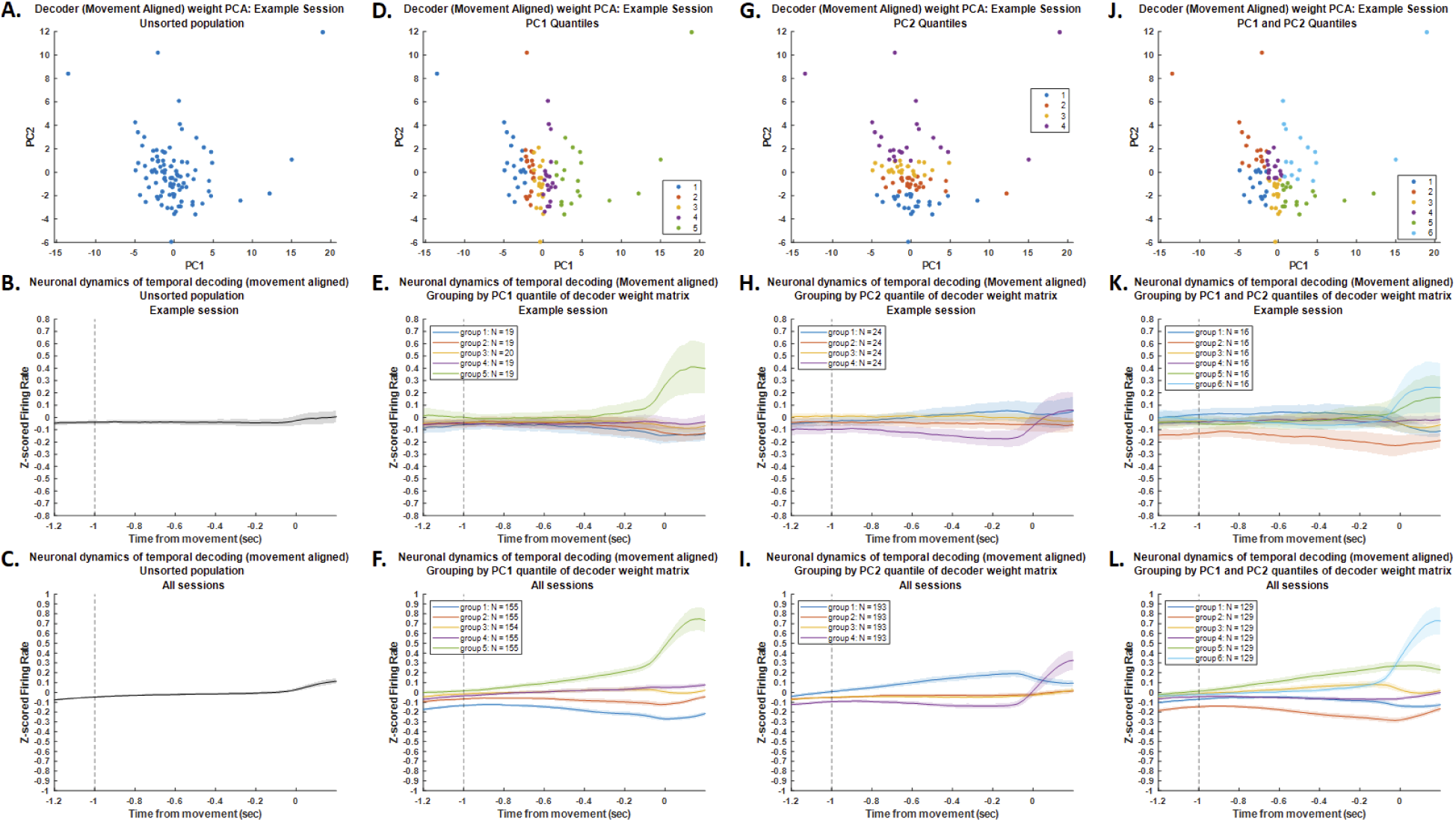
Prospective temporal encoding to movement initiation in FOF. **a**, Example session showing each unit plotted in the space of the first two principal components of the decoder weights. Each point corresponds to one unit. **b**, z-scored firing rate of all units from this session aligned to movement onset. The vertical line corresponds to the end of the decoding epoch for all panels showing neural dynamics. **c**, z-scored firing rate of all units from all sessions aligned to movement onset. **d**, Same as panel **a**, but with units sorted into quintiles along the first principal component. **e**, z-scored firing rate of all units from this session sorted into quintiles along the first principal component and aligned to movement onset. **f**, z-scored firing rate of all units from all sessions sorted into quintiles along the first principal component and aligned to movement onset. **g**, Same as panel **a**, but with units sorted into quartiles along the second principal component. **h**, z-scored firing rate of all units from this session sorted into quartiles along the second principal component and aligned to movement onset. **i**, z-scored firing rate of all units from all sessions sorted into quartiles along the second principal component and aligned to movement onset. **j**, Same as panel **a**, but with units sorted into six groups along both of the first two principal components. **k**, z-scored firing rate of all units from this session sorted into six groups along both of the first two principal components and aligned to movement onset. **l**, z-scored firing rate of all units from all sessions sorted into six groups along both of the first two principal components and aligned to movement onset. N = 16 sessions.

To further investigate this apparent difference, we next examined the “loadings” across time applied to these principal components in the PCA analysis described above. When the principal components correspond to a low-dimensional representation of weights applied to the population activity, the loadings correspond to how those principal components themselves are weighted as a function of time for our decoder. For both regions and alignments, the loading of the first principal component changed monotonically as a function of time, while the loading of the second principal component exhibited a non-monotonic trajectory, increasing to a peak before subsequently declining (**Figure 9**). This analysis was also consistent with observations of the previous analysis that ADS and FOF shared strong similarities in dynamics when aligned to stimulus start (**Figure 9a-b**). Also consistent with observations of the previous analysis is that ADS and FOF seemed to have important differences in dynamics near the time of movement initiation (**Figure 9c-d**). In particular, there is a more dramatic change in the loadings in FOF at that time compared to ADS. This was readily apparent when directly comparing the loading trajectories for matched principal components between FOF and ADS (**Figure 10**). Loading trajectories were remarkably similar between regions for both the first and second principal component when aligned to stimulus start (**Figure 10a-b**). In contrast, loadings trajectories diverged between regions when aligned to movement initiation (**Figure 10c-d**). This was present at multiple time points for the second principal component and present for both principal components leading up to the time of the movement. This suggests that ADS and FOF may have important differences that manifest most strongly near the time of the decision commitment movement.

**Figure 9.**
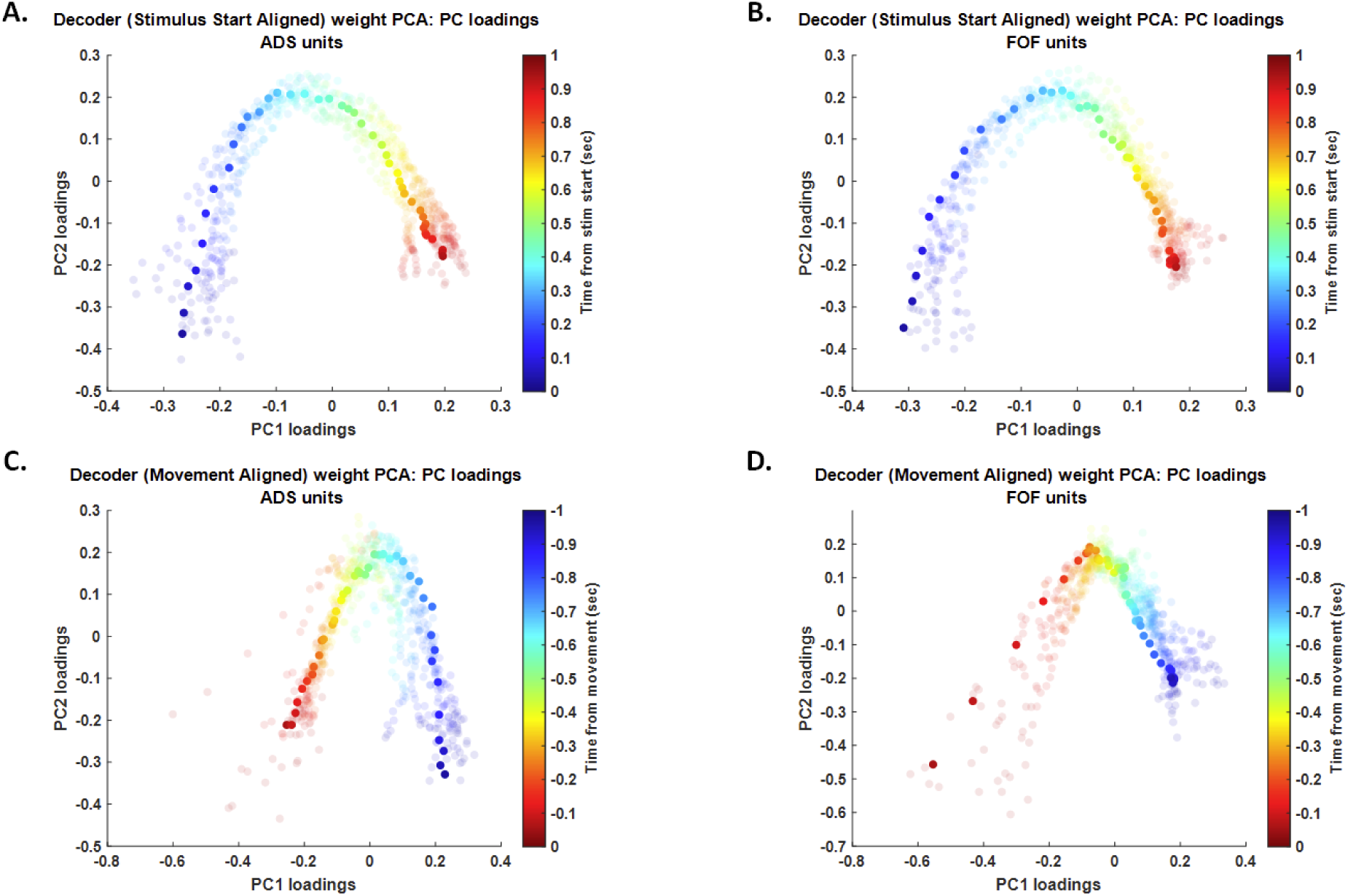
Temporal trajectory of PCA loadings from decoder weights. **a**, PCA loadings as a function of time in ADS for each session. Color indicates time from stimulus start. **b**, PCA loadings as a function of time in FOF for each session. Color indicates time from stimulus start. **c**, PCA loadings as a function of time in ADS for each session. Color indicates time from movement onset. **d**, PCA loadings as a function of time in FOF for each session. Color indicates time from movement onset. In all plots, darker points correspond to the loadings for the example session from earlier figures. N = 16 sessions.

**Figure 10.**
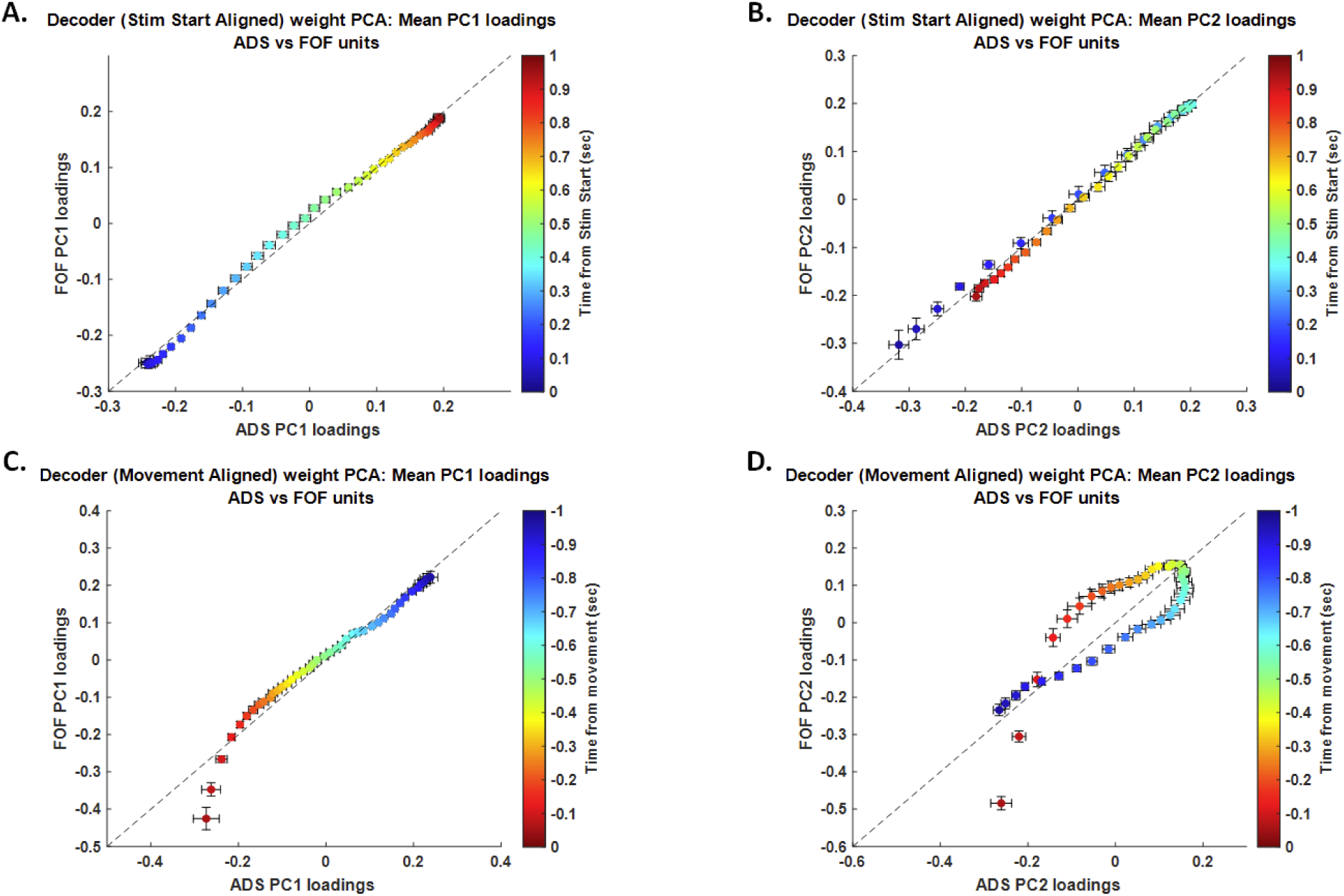
Regional comparisons of mean PCA loading trajectories across time. **a**, Comparison of loading on PC1 for FOF versus ADS aligned to stimulus start. **b**, Comparison of loading on PC2 for FOF versus ADS aligned to stimulus start. For both panels **a** and **b**, color of each point corresponds to time from stimulus start. **c**, Comparison of loading on PC1 for FOF versus ADS aligned to movement onset. **d**, Comparison of loading on PC2 for FOF versus ADS aligned to movement onset. For both panels **c** and **d**, color of each point corresponds to time from movement onset. For all plots, error bars show SEM. N = 16 sessions.

### Population Geometry

To further investigate the structure of neural trajectories across task epochs and contrast them across areas, we next turned to characterizing their population geometry. We chose a locally-linear approach, fitting separate linear models to individual task epochs, thus seeking to capture and contrast subspaces of high variance at different points in the task (see schematic in **Figure 11**,**12a**). Such an approach preserves the interpretability and robustness of linear methods with the ability to capture nonlinear dynamics (Koay et al., 2022).

**Figure 11.**
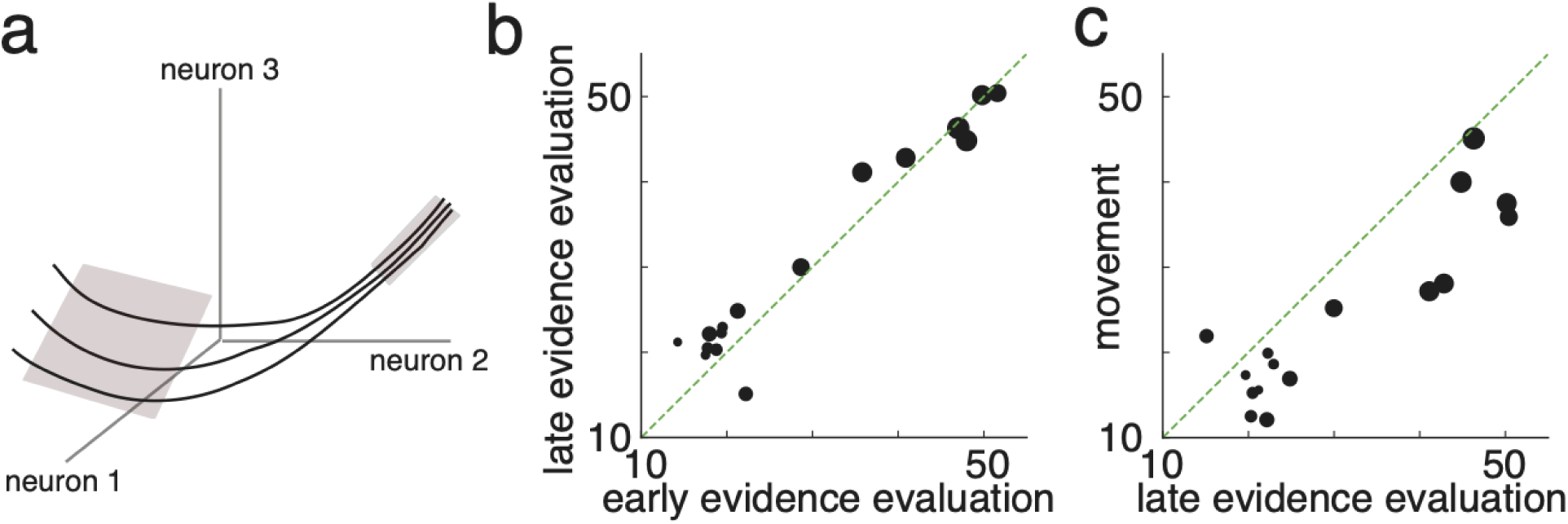
Dimension of neural population activity across trial. **a**, Schematic of locally-linear approach to characterizing dimension. PCA is used to find subspaces of high variance at different points in the task. The dimension is then calculated using the participation ratio, which measures the effective number of modes needed to capture variance. Schematic captures activity becoming lower-dimensional over the course of trial. **b**, Dimension of activity in late evidence evaluation against dimension in early evaluation, capturing stability of subspaces. **c**, Dimension of activity during decision commitment movement initiation against dimension during late evidence evaluation, capturing decrease of dimension. Sizes of points show numbers of units in each recording session.

**Figure 12.**
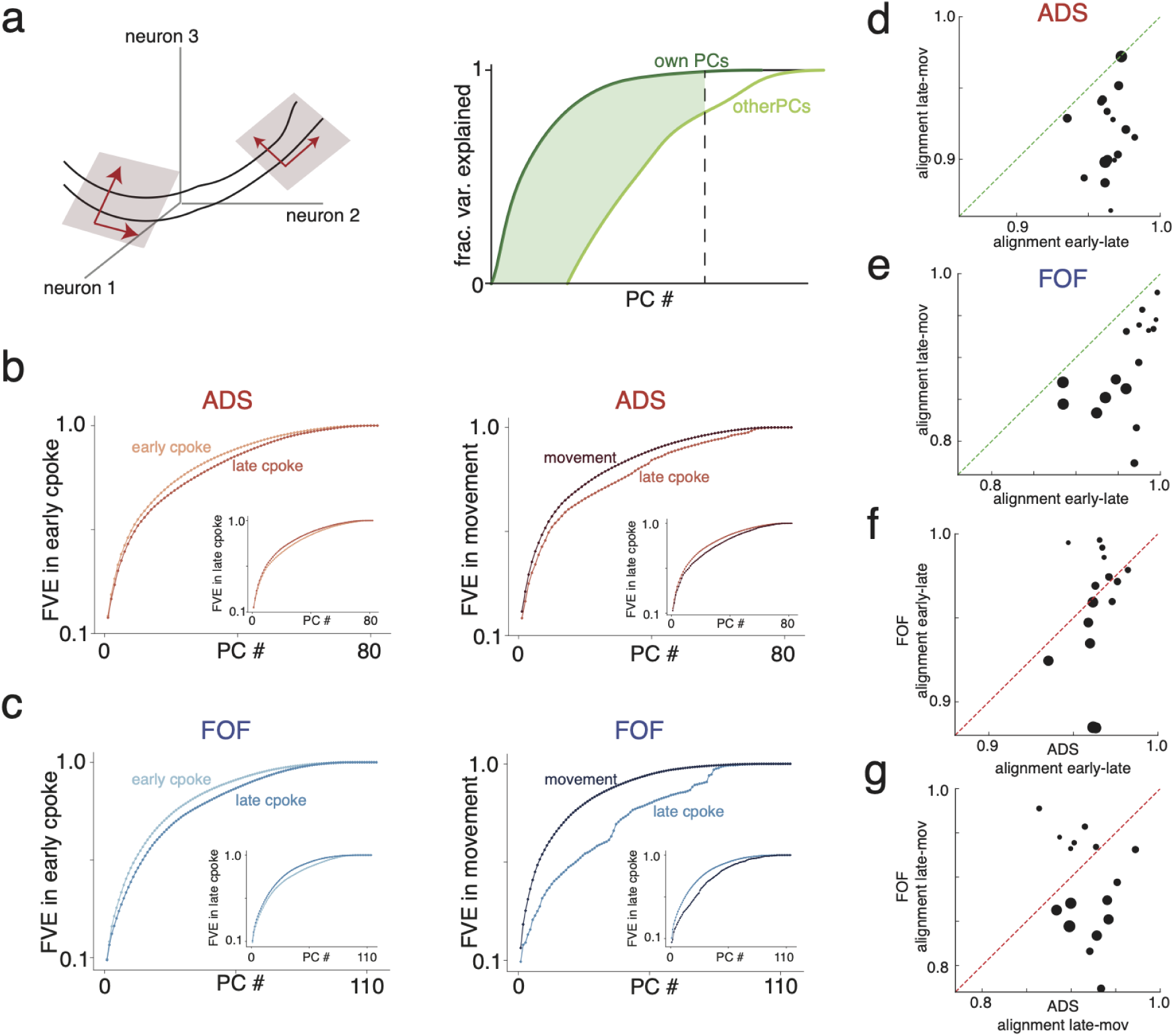
Geometry of neural population activity across trial. **a**, Left: Schematic of locally-linear approach to characterizing geometry via the alignment of subspaces of high variance at different points in the trial. Schematic shows the subspace of high variance rotating over time. Right: An alignment index is calculated by comparing how well principal component vectors (PCs) from one epoch capture population activity in another epoch when compared to PCs from that epoch. The alignment index is the area of the gap between the cumulative fraction of variance explained by each set of PCs (shared area) normalized by the total area under the higher curve (see main text and Methods for more details). **b**, Left: Cumulative fraction of variance of ADS activity during early evidence evaluation explained by early evidence PCs (top curve) and by late evidence evaluation PCs (bottom curve) for an example session. Inset shows similar plot but for activity during late evidence evaluation. Right: As in the left plot but showing movement vs. late evidence evaluation. **c**, As in (b) but for FOF activity. **d**, Scatter plot of alignment index of ADS activity between late evidence evaluation and movement against alignment index between early evidence evaluation and late evidence evaluation, showing significantly larger change upon decision commitment movement. **e**, As in (d) but for FOF activity. **f**, Scatter plot of alignment between early evidence evaluation and late evidence evaluation in FOF against corresponding alignment for ADS, showing that subspaces are similarly stable across evidence evaluation. **g**, Scatter plot of alignment between late evidence evaluation and movement in FOF against corresponding alignment for ADS, showing that change in FOF is larger. Sizes of points show numbers of units in each recording session.

We extracted subspaces of high variance by performing PCA on an epoch-specific basis, using three epochs, spanning the first 500ms of the stimulus, the last 500ms of the stimulus before the decision commitment movement, and the first 500ms upon the decision commitment movement.

To then measure and compare the dimensionality of neural population activity at different points in the task, we computed the Participation Ratio (PR) of the distribution of the explained variances. The PR captures the effective number of principal components that explain the variance in the data and is a commonly used linear metric for dimensionality (Abbott et al., 2011; Litwin-Kumar et al., 2017). We found that the dimension of neural activity remains stable across the evidence evaluation period (difference not significant, p = 0.23; **Figure 11b**), before significantly declining upon the decision commitment movement (p < 0.001; **Figure 11c**).

We next sought to capture how the subspaces of high variance changed over the course of the trial. To do this comparison, we developed an alignment index that captures the orientation of subspaces across epochs, extending an approach initiated in previous work (see Methods for a comparison of our approach with this previous approach) (Elsayed et al., 2016). The principal component vectors (PCs) derived from the neural population activity in an epoch form an ordered set of directions that best capture the variance during that epoch—the first PC captures the direction of most variance, the top 2 PCs capture the 2-dimensional plane of most variance, and so on. By contrast, PCs derived from another epoch are optimized for variance in that other epoch. If the subspaces of high variance are identical across epochs, then the second set of PCs will perform equally well as the first set of PCs when capturing variance during the first epoch. In general, however, they will do worse, and the gap in performance measures how aligned the subspaces are, while accounting for the differing importance of higher and lower variance directions. We thus computed the normalized gap in variance explained as the number of PCs increases as a measure of alignment (see schematic in **Figure 12a**, example sessions in **Figure 12b-c** and further details in Methods). When comparing two epochs, this gap can be computed using the activity in the first epoch or the activity in the second epoch (e.g., compare larger panel and inset in **Figures 12b-c**). The measures are not symmetrical and can be quite different if the activity in one epoch lies in a lower-dimensional subspace of the activity in the second epoch. Thus, to capture genuinely different subspaces, we defined our alignment index as the minimum gap across both epochs.

Despite time-varying dynamics over the evidence evaluation period, we found that the subspaces of population activity remained comparatively stable over evidence evaluation before rotating significantly upon the decision commitment movement (p < 0.001 for both areas; **Figure 12d-e**). In contrast to the general tight coordination between ADS and FOF over evidence evaluation (change in subspace alignment not significantly different, p = 0.086; **Figure 12f**), we found that the change in subspace was significantly greater in FOF than ADS (p = 0.003, **Figure 12g**). This greater change in FOF is compatible with a greater role for FOF in decision commitment.

## Discussion

The goal of this study was to understand the shared and distinct dynamical structure of frontal and striatal populations during decision making. Using large-scale Neuropixels recordings during a free-response auditory change detection task in rats, we found that both the FOF and ADS robustly encoded temporal information during decision epochs. Moreover, temporal encoding in both regions could be decomposed into two major dynamical motifs – a monotonic ramp-like mode and a transient bump-like mode. However, while these motifs were highly similar in both regions during evidence evaluation, they diverged near the time of decision commitment. FOF exhibited a sharper and more pronounced transition in neural state than ADS. This divergence was also reflected in the geometry of population activity, with FOF undergoing a significantly larger subspace change at the time of the choice report. These findings reveal a shared temporal structure across frontal and striatal circuits during deliberation and a frontal-specific reorganization that may signal decision commitment.

Our temporal decoding analyses demonstrate that both FOF and ADS contained sufficient information to recover retrospective elapsed time from stimulus onset and prospective time until movement initiation. This aligns with other work suggesting that frontal and striatal circuits maintain evolving internal estimates of time that support flexible action timing and decision control (Gouvêa et al., 2015; Jin et al., 2009; Kim et al., 2013; Mello et al., 2015; Wang et al., 2018; Xu et al., 2014; Zhou et al., 2020). The presence of matched temporal structure in both regions may result from their high interconnectivity (Cheatwood et al., 2003; Haber et al., 2000; Reep et al., 2003; Reep and Corwin, 1999), and suggests coordinated shared computational contributions.

A key finding of this study is that the dynamics in FOF and ADS began to diverge near the time of decision commitment and movement initiation. Prospective time decoding was more accurate in FOF than ADS in the final hundreds of milliseconds before the decision report, and the population geometry analyses showed that FOF underwent a sharper subspace transition at commitment. These results suggest that while both regions exhibit dynamics that could support temporal tracking during evidence evaluation, FOF may play a more active or initiating role in transitioning from deliberation to action. This aligns with prior work linking FOF to decision commitment (Hanks et al., 2015), and ADS to evidence evaluation (Yartsev et al., 2018). This also is consistent with models in which PFC initiates categorical state transitions (Hanks et al., 2015; Inagaki et al., 2022, 2019; Suzuki and Gottlieb, 2013).

Our analyses suggest that during this task frontal and striatal population activity shows highly dynamic internally-generated trajectories. We thus characterized the geometry of these trajectories with an approach adapted to such dynamic encoding, using a set of linear models fit to different task epochs. Such locally linear approaches combine the efficiency, interpretability, and robustness of linear methods with sufficient flexibility to capture highly nonlinear dynamics, and are particularly well-suited to sequential encoding of information (Koay et al., 2022). To compare the orientation of these subspaces over time, we extended a method previously proposed to analyze information encoding in the motor cortex by comparing the directions of high variance between task epochs (Elsayed et al., 2016). Our observation of a partial rotation of the subspace of high variance echoes observations from the motor cortex (Churchland and Shenoy, 2024; Elsayed et al., 2016; Kaufman et al., 2014), and may suggest a more general principle in which information that has different behavioral consequences is organized into different subspaces, allowing for differential readout and gating.

This work also naturally raises important questions about what role these temporal signals are playing in FOF and ADS and how they relate to other aspects of the decision process, such as evidence evaluation and accumulation of evidence over time. For the former, perturbation studies during task performance will be greatly beneficial to assess necessity and causal contributions. For the latter, linking the temporal motifs identified here to neural representations of evidence strength could identify how temporal and decision variables jointly shape neural trajectories. Simultaneous analysis of within- and between-area coupling and correlational structure may also provide insight into how information flows between these circuits as decisions unfold. Finally, computational modeling of recurrent population dynamics may reveal mechanisms by which these dynamical properties interact to support temporal processing and state transitions.

## Methods

### Apparatus

Tasks were programmed and run in MATLAB (Mathworks) using Bpod (Sanworks) for real-time control and measurement of behavioral output. Operant chambers used for behavioral data collection consisted of three ports. Each port contained an infrared LED beam that detects rat nose insertion upon obstruction of the beam, as well as an LED light that could be used as a cue for the rats. In addition, speakers were mounted above the left and right ports.

### Behavior

We trained rats to perform an auditory change detection task previously employed in studies involving in rats (Ganupuru et al., 2025) and human subjects (Booras et al., 2021; Ganupuru et al., 2019; Harun et al., 2020; Johnson et al., 2017). In this study’s implementation, rats inserted their nose into the central port of the behavioral apparatus cued by an LED light. Upon nose insertion, a stream of auditory pulses (“clicks”) was generated at a baseline rate of 20 or 50 Hz according to a Poisson process. Clicks were broadband and each had a duration of 3 ms. At a random point in time sampled from a truncated exponential distribution, with a minimum of 0.5 s, a mean of 2 s, and a maximum time of 4 s, the generative click rate increased by a variable magnitude of 10 Hz, 23 Hz, 36 Hz, or 49 Hz, sampled evenly. The hazard rate for change times was flat. After a change occurred, the rat had either 0.3 or 0.8 s to respond by withdrawing from the port. Successful withdrawal within the allotted time window was recorded as a “hit”, which was rewarded with a drop of water from a port on either the left or right of the central port. Failure to withdraw within the allotted time was recorded as a “miss” with reward withheld. Premature response in the absence of a change was recorded as a “false alarm” with reward withheld. Finally, on 25% of trials, no change occurred, in which case the rat was required to maintain fixation in the central port until the stimulus ended to achieve a “correct rejection”, which was rewarded. Sessions lasted for approximately 80 minutes. For behavioral criteria, we chose rats for surgical implants (see below) that had at least a 0.4 correct rejection rate and 0.5 hit rate on >300 trials per session.

### Electrophysiological recordings

Neural measurements were recorded with Neuropixel 1.0 probes (Jun et al., 2017; Steinmetz et al., 2018). Probe implants were assembled according to previous methods (Ganupuru et al., 2025; Juavinett et al., 2019). A silver wire was soldered to the grounding contacts on the probe, and the probe was glued to a 3D-printed internal mount that was affixed to a stereotaxic adapter for implantation. The internal mount was then bound to an external mount by epoxy which additionally holds a headstage circuit board for data transmission.

Probes were stereotactically implanted between +1.92 AP relative to Bregma, +1.50 mm ML relative to bregma (probe implanted contralateral to the side of the reward port assigned to the subject), and 5.50 to 6.57 mm ventral to brain surface for the probe tip. Probes were lowered slowly, roughly 20 seconds per 0.1 mm, while simultaneously recording from them to ensure probe function and verify implantation depth. Ground wires were inserted directly into the cerebellum approximately 2 mm posterior to interaural zero (“IA0”). After implantation of probe and ground wire, both craniotomies were filled with sterile optical lubricant. Absolute Dentin (Parkell) was applied to the external mount of the probe and the ground wire to bind the implant to the skull, and dental acrylic was applied over the skull to seal the implant. The headstage was taped to the external mount with Kapton tape and the implant was covered with self-adhesive wrapping. Rats were left with free access to food and water to recover for one week following surgery before resumption of training and recordings.

During recording sessions, the self-adhesive wrapping was removed from the implant, and an interface cable was plugged into the headstage. The interface cable was wrapped around a gel toe sleeve to relieve strain from the rat moving. The cable was fed through a simple pulley system to prevent the rat from grasping the cable while rearing, connecting to a PXIe acquisition module (National Instruments). This module interfaced with the Bpod and recording computer for temporal synchronization. Neural recordings were acquired using Open Ephys 3.

### Behavioral data analysis

False alarm trials included all trials in which rats withdrew their nose during stimulus baseline, including change and catch trials. Hit rates were calculated as the proportion of trials in which rats withdrew their nose within the response window (0.8 or 0.3 sec) of the change point when a change actually occurred (i.e., excluding false alarm and catch trials). Reaction times for hit trials were calculated as the time of nose withdrawal after the change point. The influence of change magnitude on hit rates was quantified via logistic regression, and the influence of change magnitude on reaction times via linear regression with p-values derived from t-statistics applied to the fitted coefficients using the Matlab GLMFIT function.

Psychophysical reverse correlations (PRCs) were generated by aligning the click times of each false alarm trial to the time of the rat’s nose withdrawal. False alarm trials were used for PRCs to avoid the confound of changes in the generative click rate. For plotting the PRCs, each false alarm click time vector was convolved with a causal half-Gaussian filter with a standard deviation of 0.05 s and sampling every 0.01 s. Regardless of false alarm time and thus duration of stimulus preceding the detection report, all false alarms were included in the PRC, with false alarms with shorter stimulus durations simply contributing to a smaller epoch of the PRC. After this smoothing, false alarm reverse correlations were averaged together. Mean increases of the click rate were calculated over the interval from 50 to 350 ms before false alarms based on the raw click counts for each trial, only including trials that were at least 350 ms in duration.

### Electrophysiological data processing and analysis

We used the Kilosort 2.5 spike sorting algorithm (Pachitariu et al., 2016) to identify single and multi unit clusters of spiking activity in our data to include for data processing. Following initial sorting, the Phy 2.0 cluster viewing software was used to manually curate clusters to remove units with drop-out due to drift via visualization of amplitude plots over the course of a session. Phy also allowed us to merge clusters that clearly originated from the same unit by assessing correlation of spike times between and within clusters and correlations in drift. Units that drifted out of the probe’s recording range during a session were excluded from analysis to avoid distortion of trial-by-trial response observations.

We further selected units for inclusion in analysis by identifying units with at least a 1 Hz firing rate during center port fixation. Peri-stimulus time histograms (PSTHs) were calculated by aligning spikes to one of two stimulus events: time of the stimulus start or movement initiation. In both cases, spikes were convolved with a Gaussian filter (0.1 s standard deviation).

### Temporal decoding of elapsed time using neuronal responses

To decode elapsed time in a trial, we leveraged Neuropixel probes to record a relatively large number of neurons in each region compared to traditional, multi-site silicon probes. In each rat for each session, we recorded from a population of neurons in FOF and ADS simultaneously. Following spike sorting, only those units that met the minimum activity threshold (see above) were used in the subsequent analyses. For each trial, spike times were aligned to the time point of interest (i.e. stimulus start or movement initiation), broken into 25ms bins, and then convolved with a causal, half-gaussian kernel (sd = 0.1 sec, sampling every 0.025 sec) to generate filtered firing rates. To decode elapsed time, the filtered firing rates served as inputs for a linear discriminant analysis (LDA) classifier, using the MATLAB “fitcdiscr” function. Elapsed time was labeled in increments of 25ms (i.e. the bin width of the filtered firing rates), relative to the time point of interest. To ensure that the analysis didn’t favor a given time bin by virtue of its prevalence, we enforced a uniform distribution of class priors by excluding all trials shorter than 1 second. This duration was calculated relative to the time point of interest while ensuring that the decoded epoch did not include times prior to the stimulus onset or following movement initiation. For stimulus-start aligned decoding, this restricted us to trials with at least 1 second between stimulus start and movement initiation or a change in the generative click rate. The decoding was then performed on this subset of trials for the 1 second following stimulus onset. For movement aligned decoding, this restricted us to trials with at least 1 second between movement initiation and stimulus start, regardless of when a change in the generative click rate occurred. The decoding was performed on this subset of trials for the 1 second prior to movement initiation. Thus, for both sets of analyses, we used the population responses in each region to decode 1 second of elapsed time, in 25ms bins, aligned to either stimulus onset (0 to 1000 ms following onset) or movement initiation (-1000 ms to 0 ms prior to movement initiation). This translated into a set of 41 classes (i.e. time bins) for each decoding analysis, as time t = 0 ms was included in each.

To account for units that varied in their range of firing rates, feature scaling was performed on the filtered firing rates. While this step isn’t necessary to assess decoder accuracy (as LDA classification maximizes class separability and exhibits invariance to scaling), the resultant decoder model coefficients (i.e. weights) are sensitive to the relative scales of features in the decoder inputs. Since we sought to use the decoder weights to classify/cluster units in each region based on their specific contributions to decoding elapsed time (see “Classifying neuronal subpopulations based on contributions to temporal decoding” below), feature scaling was necessary to ensure that the resultant decoder weights could be compared across units regardless of the range of firing rates they exhibited. We scaled decoder inputs via z-score standardization: separately for each unit, we calculated the mean and standard deviation of the filtered firing rates across all time bins of each analyzed trial (see above for inclusion criteria). These values were then used to z-score the filtered firing rate of each time bin of each analyzed trial. The z-scored filtered firing rates were then used as the input for the decoder.

Decoder performance was assessed via “leave one trial out” (LOTO) cross-validation as follows: For each analyzed trial of a given behavioral session, that trial was held out while the z-scored filtered firing rates of each unit for the remaining trials (and their associated labeled time bins) were used as the training set. We tested the held-out trial using the MATLAB “predict” function. While the training and test sets each had a uniform distribution of class priors (i.e. each time bin was equally represented in each set), we enforced no requirement that the model only predict a given time bin once per trial. As such, despite the fact that the test set (held out trial) only contained 1 of each time bin, the predict function could assign any label to any bin of the held-out trial. This ensured that each bin of the test set was assessed independently with no information about its identity gleaned from the remaining test set bins aside from their exclusion from the model’s training set.

Decoding performance was analyzed qualitatively via confusion matrices comparing the distribution of predicted vs true time bin identities for a given session and brain region (Figures 3,4 A,B). We quantified overall decoding accuracy as the root mean squared error (RMSE), in seconds. We calculated this value both across the whole session (i.e. all squared errors across all analyzed trials, averaged together); Figures 3,4 C,F) and separated by time bin (Figures 3,4 D,E). To quantify the effect of both number of units (Figures 3C,4C) and time bin (Figures 3,4 D,E) on decoder performance (RMSE), we fit the independent variables of time or number of recorded units to decoder performance via linear regression using the “fitlm” function in MATLAB. Since we observed a strong effect of the number of units on decoder accuracy (with ADS nearly always yielding more units in a given recording), we restricted our analyses comparing decoder accuracy between regions to only those sessions in which the proportion of units in FOF vs ADS was 40% or greater.

In order to compare overall decoding performance between regions we performed paired t-tests comparing the whole session RMSE for each region (Figures 3F and 4F). To quantify the difference in decoding accuracy between regions for a specific time bin, we performed bootstrap resampling of the difference in RMSE as follows. For a given temporal decoding alignment (i.e. stimulus start or movement aligned), all sessions which met our inclusion criteria for proportion of unit numbers were combined into a meta session with observed squared errors (SEs) separated by time bin. For a given time bin (e.g. 300 ms following stimulus onset, 225ms prior to movement, etc.), we calculated the difference in RMSE for the decoder constructed using ADS units vs FOF units across the entire meta session. We then compared that RMSE difference to that calculated via resampling: the SEs for each trial for the analyzed time bin from each region were shuffled together into a common pool. We then drew a bootstrap sample (with replacement) of t=number of trials in the meta session from that pool, randomly assigned them to each region, and calculated the RMSE difference as before. We repeated this 1000 times and compared the number of times the bootstrapped difference between the regions was as large or larger than the original. The proportion of repetitions that yielded RMSE differences at least as large as the observed, true RMSE difference for that time bin was used as the p-value for assessing statistical significance.

### Classifying neuronal subpopulations based on contributions to temporal decoding

We developed an analytical framework to group units based on their LDA model coefficients for classifying different time bins using dimensionality reduction via PCA. To classify units in this manner, we executed the following protocol for each analyzed session: First, LDA models were constructed for each region as before, but with all trials that met the inclusion criteria (i.e. no held out trials for cross-validation). Next, we leveraged the characteristics of a linear (as opposed to non-linear) discriminant analysis to estimate the discriminant function separately for each class (time bin). Specifically, because the covariance matrix is pooled and class independent, we can use the mean of a given class to calculate the weight vector of the discriminant function for that class (k) as follows:

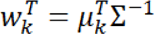

Where 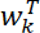 is the N units length weight vector for class k (i.e. time bin), 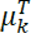 is the N units length vector of means for class k, and Σ^−1^ is the inverse of the pooled covariance matrix that is shared between all classes. Since the values (i.e. firing rates) of each class are standardized via z-scoring (see Temporal decoding), the values of that class’s corresponding discriminant function weight vector are normalized to the same range and can thus be compared across units. Concatenating the weight vectors of each class together yields an N units by T time bins matrix of discriminant function model weights representing each neuron’s relative contribution to classifying each time bin. That is, units with the highest absolute weight values in the model for a given time bin will drive the largest changes in values in classifying an unknown data point (population firing rate) as belonging to that time bin. While one could then classify units based on the time bin containing the largest weight value, this precludes classifying units as important for decoding multiple different time bins. To overcome this problem and understand how a given neuron contributes to time decoding across an entire epoch, we reduced the dimensionality of the N neuron by T time bin weight matrix using PCA (via the “pca” function in MATLAB) and classified units using the first two principal components (PCs). Overall, the first two PCs accounted for between 62% and 97% of the variance in the weight matrix for each region in a given session, making them excellent candidates for use as a basis for classifying units within a population based on their relative contribution to time decoding in different contexts.

We classified units in 3 different ways using this new basis: using even quintiles of their PC1 values (5 groups; Figures 5–8, panel D), using even quartiles of their PC2 values (4 groups; Figures 5–8, panel G), and using a combination of quantiles of both PC1 and PC2 values (3 quantiles of PC1 values split into 2 quantiles of PC2 values, for a total of 6 groups; Figures 5–8, panel J). Smoothed firing rate plots (mean +/- SEM; Figures 5–8, middle and bottom rows) for each of the groups formed by the corresponding clustering regime were calculated from the mean, normalized firing rate of each member unit. Normalization was performed by z-scoring the mean firing rate of each unit by the firing rate it exhibited during the 100 ms centered on stimulus start.

To facilitate comparison between the neuronal dynamics of each group when aligned to either stimulus start or movement onset, firing rate plots were normalized to the same point for both alignments. For the population dynamics of each group across sessions (Figures 5–8, bottom row), all data from the 16 analyzed sessions were combined into a meta-session with groups delineated by quantiles of the PC1 and/or PC2 scores as above, but for the meta-population.

### Temporal trajectory of PC loadings

In the preceding analysis, we classified units based on the values (i.e. component scores) of the first two PCs generated by a PCA designed to reduce the dimensionality of an N units by T time bins matrix of decoder weights into a 2D basis that captured most of the variance in the weight matrix. In this manner, each neuron’s vector of 41 decoder weights (one per 25ms time bin) could be expressed as 2 numbers to easily visualize and cluster their respective contributions to decoding elapsed time across the entire epoch. While the clustering itself was accomplished via the scores associated with the first two PCs, we also examined the loadings associated with each PC to understand the temporal pattern of decoder weights associated with the units projected onto those PCs. That is, while clustering the population based on the first two PC scores allowed us to group units with a similar decoding weight profile together, examining the loadings of a given PC allowed us to visualize the “shape” of the decoder weight profiles associated with units that have high scores in that PC.

To visualize the trajectory of the “loadings” of the first two PCs over time for each alignment regime and analyzed region, we performed a modified rectification of the loading values output by the same “pca” MATLAB function which calculated the PC “scores” used above. The purpose of this rectification was to ensure that the “shape” of the loading value “trajectory” was oriented in the same direction for each session in order to facilitate comparison. To do this, we applied the following procedure: for a given session, if the final time bin of the epoch (i.e. 1 sec following stimulus onset or -1 sec preceding movement) had a negative PC1 loading value, change the sign of all of the PC1 loading values for that session. Similarly, if the final time bin of the epoch had a positive PC2 loading value, change the sign of all of the PC2 loading values in that session. In this manner, the magnitude of each loading value was preserved while ensuring that the loading value associated with the final time bin was plotted in the bottom right and the value associated with the first time bin (“anchor time”) was plotted in the bottom left (Figure 9).

### Population geometry

For the population geometry analyses, we retained trials where the center-poke interval was greater than 1 s. We then divided each trial into 3 epochs. The first epoch is the first 500ms following center-poke start and called “early cpoke”. The second epoch is the last 500ms of the center poke (aligned to center poke-out) called the “late cpoke”. The third epoch is the 500ms after the center poke-out called the “movement” epoch. We checked that the rat took at least 500ms to move from the center port to the reward port in all trials.

For these analyses, we only considered single units. Spikes were binned in 25 ms bins and smoothed with a causal half-Gaussian kernel of 100 ms. Thus, for each trial we have an *N* x *T* firing rate matrix of *N* neurons and *T* time steps, which is *T* = 500 ms / 25 ms = 20 time bins for the epochs described above. Correspondingly, we have a *N* x (*n_trials_T*) matrix for a single epoch of a single session. The analyses described below are performed on the firing rate matrices which we call *FR*.

The neuron-neuron covariance matrices of the *FR* matrices are defined as

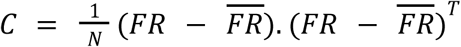

where 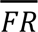 is the average firing rate of each neuron over *n_trials_T* time bins. Note that the average firing rate 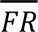 is calculated in each epoch for the *n_trials_T* time bins and then subtracted from each column of matrix *FR* in that epoch.

As described below, we use these covariance matrices to calculate measures of dimensionality and subspace alignment over the course of the trial. To compare these measures across epochs, we use weighted paired t-tests with the weight of a session given by the number of neurons recorded in that session.

### Measurement of dimension via participation ratio

The *N* eigenvalues of the *N* x *N* covariance matrix of neural activity capture the spread of the activity along various dimensions in neural state space, with bigger eigenvalues indicating more spread along the corresponding eigendimension of the covariance matrix. A single large eigenvalue and very small remaining eigenvalues indicate data that is near one-dimensional, while *N* equally-large eigenvalues indicate data that is *N*-dimensional and thus as high-dimensional as can be given the number of neurons. Correspondingly, the dimensionality of the data can be measured using the spread of the eigenvalues.

Participation ratio (PR) is a commonly used measure of the spread of the eigenvalues, and captures the effective number of dimensions that contribute to neural population activity (Abbott et al., 2011; Litwin-Kumar et al., 2017). It is defined in terms of the eigenvalues of the covariance matrix as

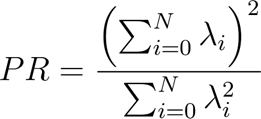

where *λ_i_* is the *i*th eigenvalue. We calculate the PR for each epoch described above for each session.

We note that measures of dimension based on the eigenvalues of the covariance matrix are linear measures, capturing the effective dimension of the linear subspace that contains most of the variance. The locally-linear approach we use involves fitting different linear models to different regions of state space and thus can account for nonlinear structure in the data.

### Measure of subspace alignment via Alignment Index

The alignment index uses Principal Components Analysis (PCA) to compare the orientation of high variance subspaces across two epochs. Let the two epochs be *E_1_* and *E_2_* respectively, with covariance matrices *C_1_* and *C_2_*. Let the columns of the *N*x*N* matrix *V_1_* correspond to the principal component vectors from epoch 1 (i.e., the eigenvectors of C_1_), ordered by the magnitude of the corresponding eigenvalue, and define *V_2_* similarly. We will also use the notation *V_1_*(:, :P) to denote the first P columns of the matrix *V_1_* (i.e., the *P* eigenvectors corresponding to the *P* largest eigenvalues). Finally, recall that the trace of a matrix *C*, Tr*(C)*, is the sum of the diagonal entries and is also equal to the sum of the eigenvalues.

Note that the total variance of neural activity during epoch *i* is Tr(*C_i_*). Moreover, the total variance of neural activity during epoch *i* when projected into the subspace spanned by the first P columns of *V_j_* is Tr(*V_j_*(:, :P)^T^ *C_i_V_j_*(:, :P)). Thus, the fraction of variance of neural activity during epoch *i* explained by the first *P* PCs in epoch *j* is

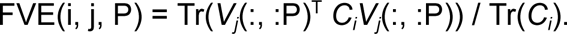

The PCs from epoch *i* are optimized to capture variance during epoch *i* and thus will always perform at least as well as (and typically better than) PCs from another epoch. Thus for any *P*, FVE(i,i,P)>=FVE(i,j,P). In previous work, the ratio of these quantities was used as a measure of alignment between population activity in different epochs, with the choice *P=10* (i.e., looking at the top 10 PCs) (Elsayed et al., 2016)

Rather than choosing a fixed *P* as a threshold, we considered the curves FVE(i,i,P) and FVE(i,j,P) as functions of *P* (see Figure 11a-c). We then calculated the approximate area between the curves from 0 to *K_i_*, the point where the top curve saturated, defined as the minimum number of PCs required to explain *95%* of the variance (i.e., *FVE(i, i, K_i_)>=0.95*). We normalized this area by the total area under the upper curve between 0 and *K_i_*. Thus,

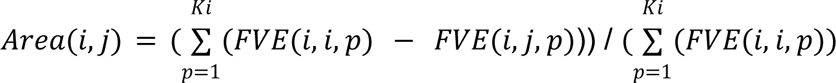

These areas may not be symmetrical. In particular, if activity during one epoch occupies a lower-dimensional subspace of activity during another epoch, then the area between one pair of curves could be small and the other large. Thus, to capture differences in overall alignment rather than subspace structure, we define the alignment index using the minimum of the areas calculated using population activity in epochs *i* and *j*. We moreover subtract it from 1 so that a smaller area between the curves leads to higher values of alignment index. This procedure yields the measure for alignment index

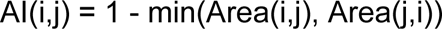

